# Spatially localized cluster solutions in inhibitory neural networks

**DOI:** 10.1101/2020.07.30.229542

**Authors:** Hwayeon Ryu, Jennifer Miller, Zeynep Teymuroglu, Xueying Wang, Victoria Booth, Sue Ann Campbell

## Abstract

Neurons in the inhibitory network of the striatum display cell assembly firing patterns which recent results suggest may consist of spatially compact neural clusters. Previous computational modeling of striatal neural networks has indicated that non-monotonic, distance-dependent coupling may promote spatially localized cluster firing. Here, we identify conditions for the existence and stability of cluster firing solutions in which clusters consist of spatially adjacent neurons in inhibitory neural networks. We consider simple non-monotonic, distance-dependent connectivity schemes in weakly coupled 1-D networks where cells make strong connections with their *k^th^* nearest neighbors on each side. Using the phase model reduction of the network system, we prove the existence of cluster solutions where neurons that are spatially close together are also synchronized in the same cluster, and find stability conditions for these solutions. Our analysis predicts the long-term behavior for networks of neurons, and we confirm our results by numerical simulations of biophysical neuron network models. Additionally, we add weaker coupling between closer neighbors as a perturbation to our network connectivity. We analyze the existence and stability of cluster solutions of the perturbed network and validate our results with numerical simulations. Our results demonstrate that an inhibitory network with non-monotonic, distance-dependent connectivity can exhibit cluster solutions where adjacent cells fire together.

## 1. Introduction

Many types of brain activity are characterized by coordinated activity of neural assemblies, in which neuron firing is synchronized within an individual assembly but not between different assemblies [18, 27, 19, 11, 32, 6]. Neural assemblies have been observed between neurons in different cortical columns [18], within regions of the hippocampus [19, 11], the dentate gyrus [32] and between cells in the striatum [6, 9, 2] and the olfactory bulb [27]. Neural assemblies may involve neurons which are widespread across one or more brain regions [18, 19] or may involve spatially localized neurons [32, 6]. Understanding the dynamics and formation of neuronal assemblies within larger neural networks has gained increasing importance in neuroscience [12] and has been studied both experimentally [27, 19, 15, 32, 6] and using computational modeling [16, 28, 17, 8, 1, 22, 3, 34].

In neural network models, the formation of neural assemblies has been analyzed by identifying cluster solutions in networks of intrinsically oscillating neurons [16, 17, 28, 15, 22, 31, 7]. Clustering defines a type of solution where the network of oscillators breaks into subgroups. Within each subgroup, the phases of the oscillators are the same, while oscillators in different subgroups are phase-locked with some nonzero phase difference. A useful mathematical framework for studying cluster solutions is the phase model reduction [21, 37]. This framework has been used to study synchronization and clustering in a variety of coupled oscillator networks [5, 33, 25, 36, 29].

In previous work [31, 7], we used the phase model approach to determine existence and stability conditions of cluster solutions of networks with neurons arranged in a 1-dimensional ring, of arbitrary size, with various connectivity schemes. As in many other studies [5, 33], our work focused on cluster solutions where the phase difference between any two adjacent neurons in the network is the same. Thus, other than the case of homogeneous all-to-all coupling, cells in the same cluster were dispersed throughout the network.

Neural assembly firing has been identified in the striatum, a sparsely connected, inhibitory network that is part of the basal ganglia circuit [30, 9, 2, 6, 23]. Recent experimental imaging of inhibitory medium spiny neuron firing in the striatum has suggested that assemblies can be spatially compact [6]. While computational modeling of inhibitory striatal networks has primarily investigated the formation of neural assemblies in which the assembly cells are spatially dispersed in the network [34, 3], a recent study found that spatially compact cluster firing patterns can result from non-monotonic, distance-dependent connectivity in which cells made stronger synaptic connections with their more spatially distant neighbors compared to their nearest neighbors [38]. This result suggests that in inhibitory networks neurons located near one another should be able to be a part of the same cluster when they are more strongly connected to neighbors farther away.

In this work, motivated by the results in [38], we study 1-D ring, inhibitory networks with simple non-monotonic, distance-dependent connectivity schemes. Specifically, we consider networks in which neurons are connected to only their *k^th^* nearest neighbors and identify conditions for the existence and stability of solutions which exhibit spatially localized cluster firing. We additionally consider connectivity schemes with connections between the first to (*k* - 1)^*th*^ nearest neighbors that are weaker than connections to *k^th^* nearest neighbors, and analyze how this additional local coupling between cells affects the existence and stability of spatially localized cluster solutions. We employ a phase model reduction of the network to obtain analytical conditions and then test the conditions with numerical simulations of biophysical neural network models.

Our paper is structured as follows: Section 2 provides the description of the methodology we employ, including the phase model reduction (Sec. 2.1) for analysis and the biophysical neuron model (Sec. 2.2) for numerical simulations. Section 3 describes our analysis and numerical results for the case of only *k^th^* nearest neighbor coupling. Section 4 presents results for the case of additional coupling(s) between the first *k* nearest neighbors. We conclude with a discussion of our results in Section 5.

## 2. Methodology

We begin by reviewing the phase reduction method that reduces a weakly coupled neural network model to a phase model. This phase model is used to conduct our analysis. We then introduce the specific neural network model that we use for numerical simulations and include the parameter values for our simulations.

### 2.1. Phase reduction method

Consider a general network model consisting of *N* identical, weakly coupled oscillators on a ring with circulant coupling

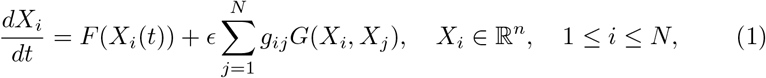

where *ϵ* is the coupling strength with 0 < *ϵ* ≪ 1, *F* is the vector field of the isolated neuron, *G* is the coupling function and *W* = (*g_ij_*) is the coupling matrix with *g_ij_* = *g*_*j*−*i*(mod *N*)_ and *g_ii_* = 0 for *i, j* = 1, 2,…, *N*.

We assume that when isolated from the network each neuron exhibits an exponentially asymptotically stable *T*-periodic orbit, denoted as 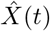, for 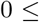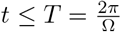, which is a solution of

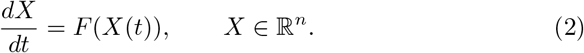

Applying the theory of weakly coupled oscillators [14, 21, 37], the complete state of each neuron in the network can be approximated by its phase on its *T*-periodic limit cycle, *θ_i_*(*t*) = Ω*t* + *ψ_i_*(*t*) ∊ [0, 2*π*), where *φ_i_*(*t*) is the relative phase of the *i^th^* neuron. Hence, this theory enables us to significantly reduce the number of equations that describe a neuronal network from *n* equations to one per neuron.

The dynamics of the relative phase of the *i^th^* neuronal oscillator is slowly varying and governed by the equation

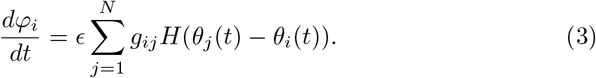

Here *H* is known as the interaction function and is given by

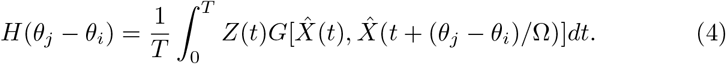

*H* captures the modulation of the instantaneous phase of the *i^th^* oscillator due to the coupling. *Z* is referred to as the phase response curve of the unperturbed oscillator, which is the unique periodic solution of the linearized adjoint system

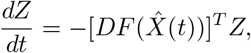

subject to the normalization condition

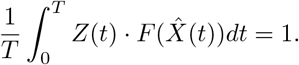

Thus, the corresponding phase model is given by

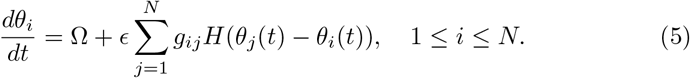

In view of circulant coupling (*g_ij_* = *g*_*j*−*i*(mod *N*)_ for 1 ≤ *i, j* ≤ *N*) and no self-coupling (*g_ii_* = *g*_0_ = 0, 1 ≤ *i* ≤ *N*), the phase model (5) can be written as

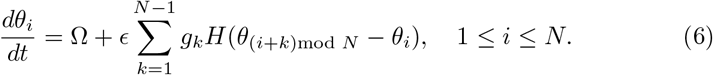

We will use this phase model to determine the existence and stability of certain cluster solutions and how this depends on the connections *g_k_*, focusing on results that can be applied to any neural model (1). We then use these results to predict which cluster solutions will be stable in the neural network model described below.

### 2.2. Neural network model

To verify our analysis results, we numerically simulate networks of neurons modeled by the conductance-based Wang and Buzsáki inhibitory interneuron model [41]. This model uses the classic Hodgkin-Huxley formalism [20] with parameters adjusted to match the action potential shape and spiking properties of fast-spiking interneurons. The membrane voltage *V* of each individual neuron is governed by the following equations:

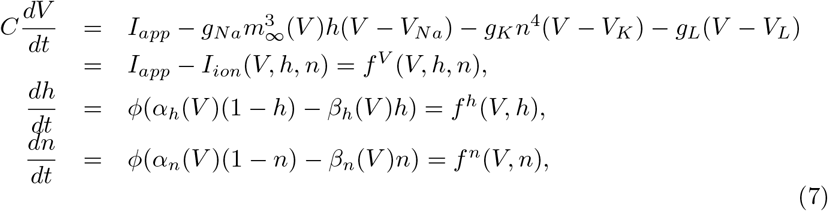

where *t* is time in mS and *V* is the cell membrane potential in mV. The variables *h* and *n* are, respectively, the inactivation gating of the sodium current and the activation gating of the potassium current. The sodium current is assumed to instantaneously activate according to the steady state activation function *m*_∞_(*V*) = *α_m_*(*V*)/(*α_m_*(*V*) + *β_m_*(*V*)), where *α_m_*(*V*) = −0.1(*V* + 35)/(exp(−0.1(*V* + 35)) − 1), *β_m_*(*V*) = 4 exp(−(*V* + 60)/18) are the voltage dependent reaction rates associated with the activation gate with units ms^−1^. The reaction rates for the inactivation of the sodium channel and activation of the potassium channel are given by: *α_h_*(*V*) = 0.07 exp(−(*V* + 58)/20), *β_h_*(*V*) = 1/(exp(−0.1(*V* + 28)) + 1), *α_n_*(*V*) = −0.01(*V* + 34)/(exp(−0.1(*V* + 34)) − 1), *β_n_*(*V*) = 0.125 exp(−(*V* + 44)/80). We model the situation where each neuron is intrinsically firing at a biologically reasonable rate of less than 60 Hz [41, 26] by setting *I_app_* < 1 *μ*A/cm^2^.

In the network, neurons are coupled with fast inhibitory synapses that are modeled using first order kinetics following [10]:

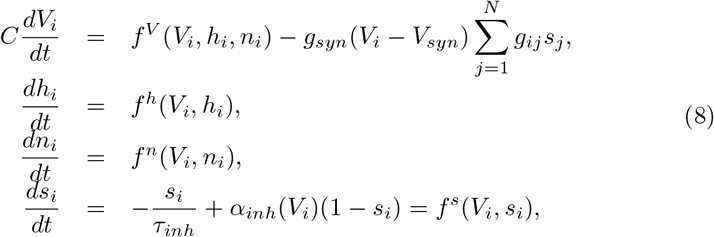

where *g_syn_* is the maximal synaptic strength and *g_ij_* scales the strength of the synaptic current from cell *j* to cell *i*. The synaptic gating variable *s_i_* for presynaptic cell *i* depends on membrane voltage *V_i_* according to

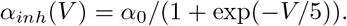

Descriptions of the parameters and values used in our numerical simulations are given in Table 1.

**Table 1:**
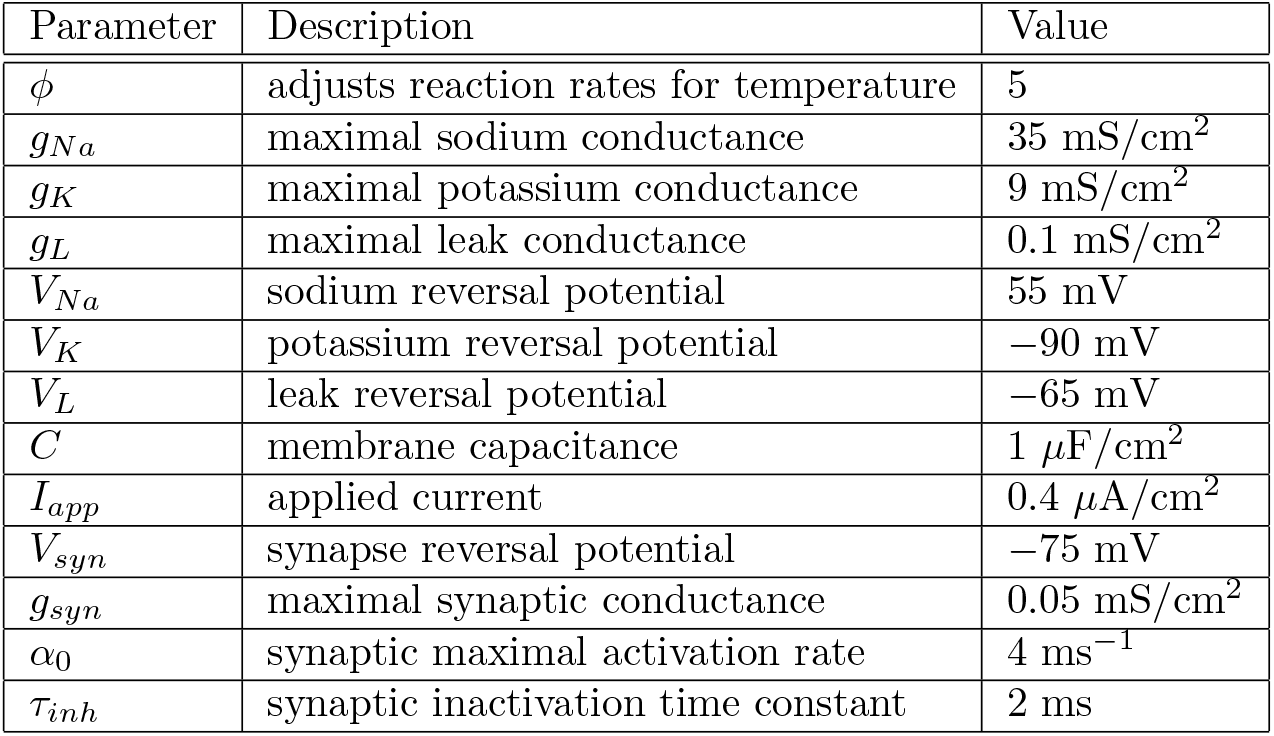
Description of parameters and the values used for the neuron model in (8).

| Parameter | Description | Value |
| --- | --- | --- |
| $\phi$ | adjusts reaction rates for temperature | 5 |
| $g_{Na}$ | maximal sodium conductance | 35 mS/cm<sup>2</sup> |
| $g_K$ | maximal potassium conductance | 9 mS/cm<sup>2</sup> |
| $g_L$ | maximal leak conductance | 0.1 mS/cm<sup>2</sup> |
| $V_{Na}$ | sodium reversal potential | 55 mV |
| $V_K$ | potassium reversal potential | −90 mV |
| $V_L$ | leak reversal potential | −65 mV |
| $C$ | membrane capacitance | 1 μF/cm<sup>2</sup> |
| $I_{app}$ | applied current | 0.4 μA/cm<sup>2</sup> |
| $V_{syn}$ | synapse reversal potential | −75 mV |
| $g_{syn}$ | maximal synaptic conductance | 0.05 mS/cm<sup>2</sup> |
| $\alpha_0$ | synaptic maximal activation rate | 4 ms<sup>−1</sup> |
| $\tau_{inh}$ | synaptic inactivation time constant | 2 ms |

Comparison of numerical results of this interneuron network model with the predictions given by the phase reduction model requires the interaction function *H* in Eq. (6). The model (8) can be put in the form given in (1) if we identify *ϵ* = *g_syn_*, *X_i_* = (*V_i_, h_i_, n_i_, s_i_*)^*T*^, *F* = (*f^V^, f^h^, f^n^, f^s^*)^*T*^ and *G*(*X_i_, X_j_*) = ((*V_syn_* − *V*_*i*_)*s_j_*, 0, 0, 0)^*T*^. Then the function *H* can be computed from (4). For any neural network, *H* rarely has a closed-form expression and one usually has to resort to numerical evaluation. In this paper, we use XPPAUT [13] to numerically compute *H* for the Wang-Buzsáki inhibitory network (8).

## 3. The First Case: *k^th^* Nearest Neighbor Coupling Only

In this section, we first study the cluster solutions of an *N*-cell network with *k^th^* nearest neighbor coupling only; i.e., *g_k_* > 0, *g_N_*_−*k*_ > 0, and *g_i_* = 0 otherwise. We look for solutions with *m* clusters. We assume that *N* = *mkp* for some 1 ≤ *p* ≤ *N*. We show two examples of this type of network solution with *N* = 12 in Figure 1. In Fig. 1(a), there are *m* = 2 clusters (cells represented by circular nodes all fire together as one cluster and those represented by rectangular nodes fire together as the second). The cells have reciprocal, second nearest neighbor coupling, *k* = 2 (e.g., cell 1 is coupled to cells 3 and 11). There are *p* = 3 subgroups in each cluster (e.g., the cluster of cells represented by circular nodes contains three spatially separated subgroups: 1 & 2, 5 & 6, and 9 & 10). The second nearest neighbor coupling causes the network to have two disjoint subnetworks, shown on the right, i.e., odd-numbered vs. even-numbered cells.

**Figure 1:**
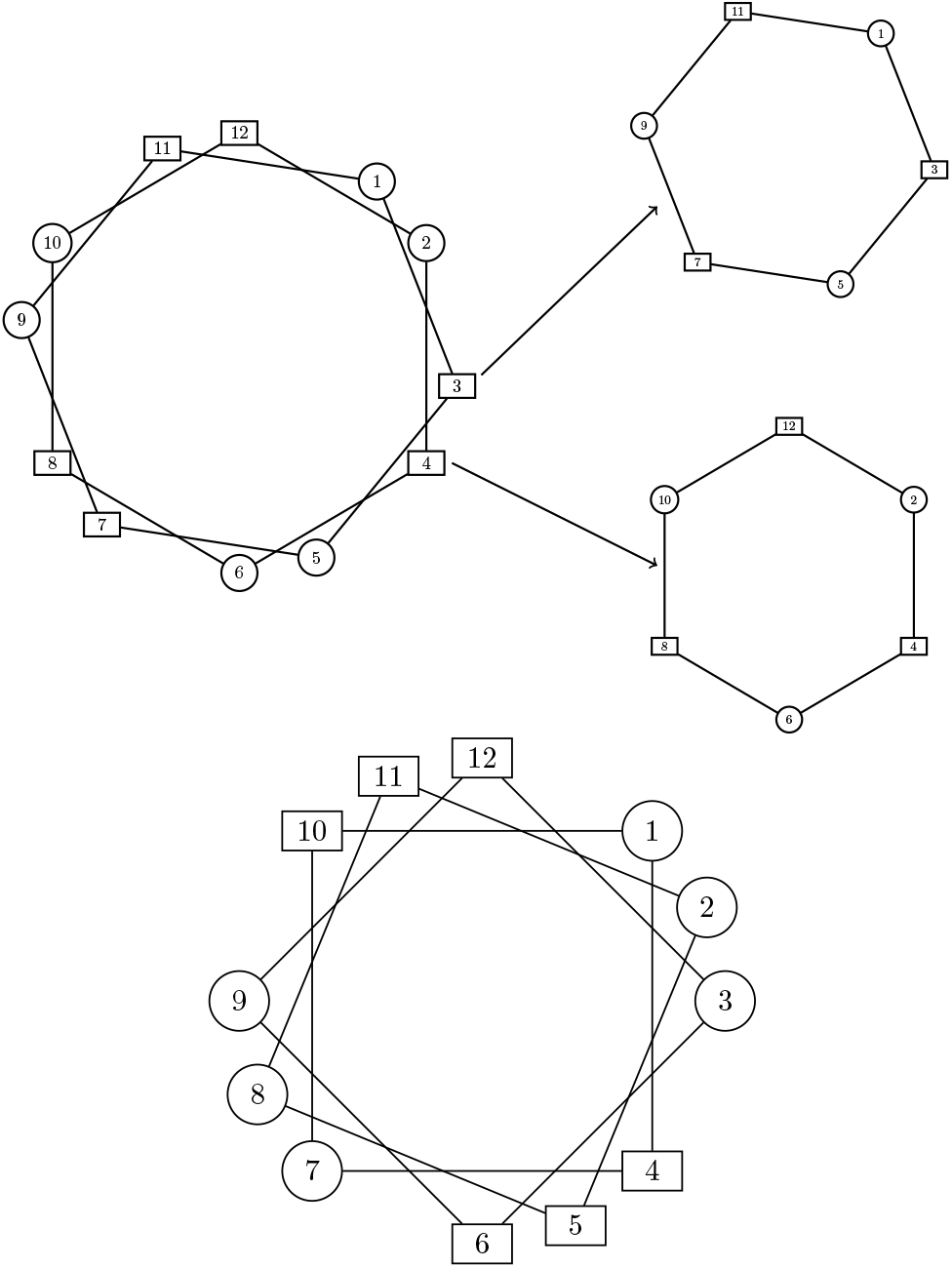
(a) Top: the full network for *N* = 12, *m* = 2, *k* = 2, *p* = 3, which can be decomposed into the two disjoint networks of 6 cells each, i.e., odd-numbered vs. even-numbered cells; (b) Bottom: the network of *N* = 12, *m* = 2, *k* = 3, *p* = 2.

Fig. 1(b) shows a network solution with the same number of cells but with different connectivity. Each neuron is only coupled to its third nearest neighbor (*k* = 3) on each side and the network of *N* = 12 decomposes into 3 disjoint sub-networks, each containing 4 neurons. In general, *k^th^* nearest neighbor coupling results in *k* disjoint subnetworks with *mp* neurons in each subnetwork.

### 3.1. Cluster Solutions

This subsection begins by showing the existence of the *m*-cluster solutions with this specific network structure and finding stability conditions. Next, we apply the results to make predictions for the behavior of examples with *N* = 8, 12, 18. Finally, we compare these predictions to numerical simulations of the example networks.

#### 3.1.1. Existence and stability

Define the phase difference *ϕ_i_* = *θ*_(*i*+1)mod *N*_ − θ_*i*_, *i* = 1,…, *N*. We prove the following theorem which shows that, for networks with the structure described above, certain *m*-cluster solutions of the type shown in Figure 1 always exist.

Further, the stability of these solutions depends on the phase difference between the clusters.

##### Theorem 3.1.

*For any given network structure with N* = *mkp neurons and g_k_* > 0, *g_N_*_−*k*_ > 0 *and g_i_* = 0 *otherwise, there exists an m-cluster solution of the following form*:

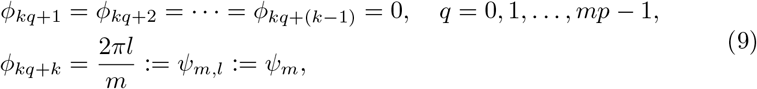

*with gcd*(*l, m*) = 1 *and l* < *m. Moreover, the m-cluster solution (9) is stable if*

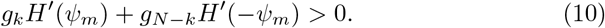

*In particular, if g_k_* = *g_N_*_−*k*_, *then this m-cluster solution is stable if H*′_*odd*_(*ψ_m_*) > 0.

*Proof.* **Existence**: We look for the solution of the form 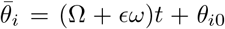 where *θ*_10_ = 0 and 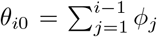 for 2 ≤ *i* ≤ *N*. If 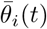 is a solution of (6), then

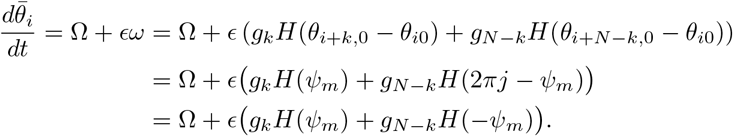

Thus, *ω* = *g_k_H*(*ψ_m_*)+*g_N_*_−*k*_*H*(−*ψ_m_*). It is clear that, in (9), 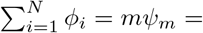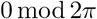, which leads to an *m*-cluster solution in the network of size *N* = *mkp*.

**Stability**: Let 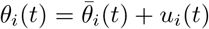. Then

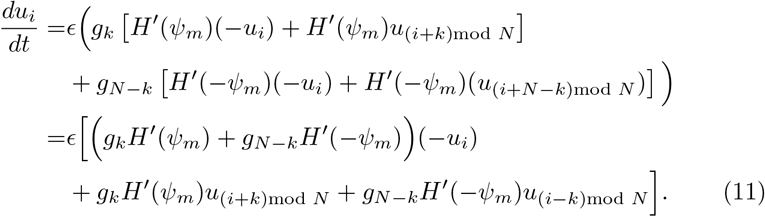

Let *A* = *circ*(*a*_1_, *a*_*2*_,…, *a_N_*) for which

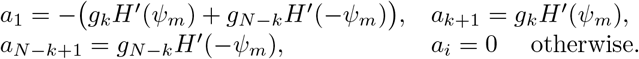

Rewriting (11) in vector form, we have

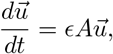

where 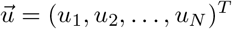. The stability of this vector equation is determined by the real part of the eigenvalues of the Jacobian matrix *A*. Let 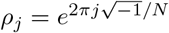 for 1 ≤ *j* ≤ *N*. Because *A* is circulant and *ρ^N^* = 1, the eigenvalues of *A* are given by

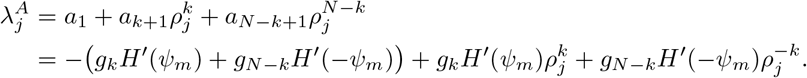

Hence, the real part of the eigenvalues is

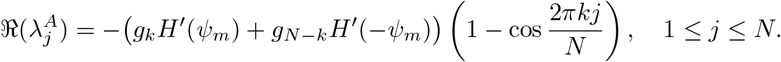

Note that 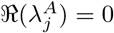 whenever 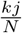 is an integer. Thus there are *k* zero eigenvalues. When *g_k_* = *g_N_*_−*k*_, then

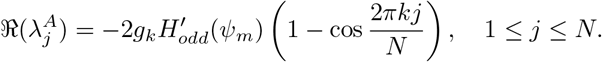

Therefore, the *m*-cluster solution is stable if

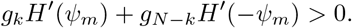

Additionally, when *g_k_* = *g_N_*_−*k*_, then this *m*-cluster solution is stable if *H′_odd_*(*ψ_m_*) > 0.

We note that the existence of *m*-cluster solutions is determined only by the network structure. Additionally, given symmetric connection weights, the stability of the solutions is independent of the network size. Due to the zero eigenvalues, we can obtain stability but not asymptotic stability of these solutions when *k >* 1. We will discuss these zero eigenvalues further in Sec. 3.2.1. When *k* = 1, however, there is only one zero eigenvalue which corresponds to motion along the solutions [7]. Thus for *k* = 1 the condition (10) does give asymptotic stability. This is consistent with the stability result in [31] where we showed that an *m*-cluster solution is asymptotically stable if *H′_odd_*(*ψ_m_*) > 0, in the case of symmetric connectivity with only first nearest neighbor coupling.

#### 3.1.2 Specific network examples

To clarify our results and for comparison with the numerical simulations in the next section, we now apply the phase model results to some examples for specific values of the total number of neurons *N*, with *N* = *mkp* for some integers *m, k, p* with *m* > 1.

Given *N* and *k*, from Theorem 3.1, the stability condition for any appropriate *m* is *g*_*k*_*H*′(−*ψ_m_*) + *g_N_*_−*k*_*H*′(*ψ_m_*) > 0 where *ψ_m_* is the phase difference between clusters as defined in the theorem. In the numerical simulations below we use symmetric coupling, *g_k_* = *g_N_*_−*k*_ > 0, so the stability condition reduces to *H′_odd_*(*ψ_m_*) > 0, where *H_odd_* is the odd part of *H*. Hence for a given interaction function *H*, stability depends only on the phase difference and not the coupling strengths. Thus to find the stability of an *m*-cluster solution, we need only calculate *ψ_m_* and determine the sign of *H′_odd_*(*ψ_m_*).

We used XPPAUT [13] to numerically compute the functions *H* and *H_odd_* for the model (8) with parameters given in Table 1. The results are shown in Figure 2. Careful inspection of the results shows that *H′_odd_*(*ϕ*) is positive for *ϕ* ∈ [0, 0.11*π*) ∪ (0.62*π*, 1.38*π*) ∪ (1.89*π*, 2*π*] and negative everywhere else.

**Figure 2:**
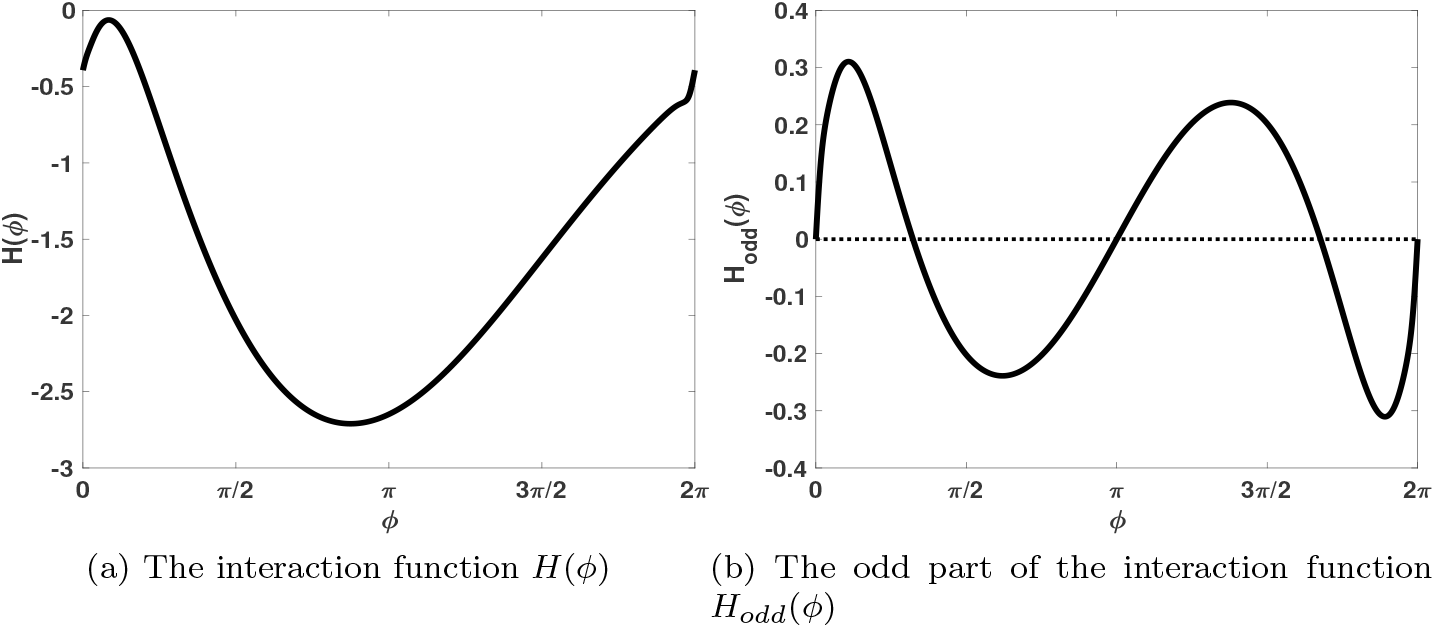
Profile information of the interaction function *H* for the neural model (8) with parameter values given in Table 1.

In [7] it is shown that the synchronous (1-cluster) solution exists for all values of *N* and any circulant coupling matrix, and that this solution is asymptotically stable if *H*′(0) > 0. Thus from Figure 2 we predict that the synchronous solution is asymptotically stable for the specific neural model we consider, given in Eq. (8), with any circulant coupling matrix.

Now consider any *N*, with *k*th nearest neighbor coupling. For *m* = 2, *ψ_m_* = *π* and *H′_odd_*(*π*) > 0 so the 2-cluster solution is predicted to be stable. For *m* = 4, *ψ_m_* = *π/*2 or *ψ_m_* = 3*π*/2. Since *H′_odd_*(*π*/2) < 0 and *H′_odd_*(3*π*/2) < 0 the 4-cluster solution is predicted to be unstable. Now with *N* = 8 and *k* = 2 one may have *m* = 2 or *m* = 4. Thus for *N* = 8 with second nearest neighbor coupling the prediction is that only the synchronous and 2-cluster solutions are stable. We summarize the predictions for other values of *N* in Table 2.

**Table 2:** Stability predictions from phase model analysis for the network (8).

| $N$ | $k$ | $m$ | $p$ | $\psi_m$ | Prediction |
| --- | --- | --- | --- | --- | --- |
| 8 | 2 | 2 | 2 | $\pi$ | stable |
| | | 4 | 1 | $\pi/2, 3\pi/2$ | unstable |
| | 4 | 2 | 1 | $\pi$ | stable |
| 12 | 2 | 2 | 3 | $\pi$ | stable |
| | | 3 | 2 | $2\pi/3, 4\pi/3$ | stable |
| | | 6 | 1 | $\pi/3, 5\pi/3$ | unstable |
| | 3 | 2 | 2 | $\pi$ | stable |
| | | 4 | 1 | $\pi/2, 3\pi/2$ | unstable |
| 18 | 2 | 3 | 3 | $2\pi/3, 4\pi/3$ | stable |
| | | 9 | 1 | $2\pi/9, 4\pi/9, 14\pi/9, 16\pi/9$ | unstable |
| | | | | $8\pi/9, 10\pi/9$ | stable |
| | 3 | 2 | 3 | $\pi$ | stable |
| | | 3 | 2 | $2\pi/3, 4\pi/3$ | stable |
| | | 6 | 1 | $\pi/3, 5\pi/3$ | unstable |

#### 3.1.3 Numerical results

We numerically simulate networks of *N* neurons (where *N* is even) with only *k^th^* nearest neighbor coupling to validate the existence and stability of *m*-cluster solutions, as discussed in Sec. 3.1.1. For all simulations, we use the model in (8) and the parameter values given in Table 1, including synaptic coupling strength *g_syn_*, implemented in Matlab. Only the number of cells in the network *N* and the connectivity pattern (second or third nearest neighbor coupling) are varied.

We first consider the case for second nearest neighbor coupling only, i.e., *k* = 2. We use random initial conditions and simulate at least up to *t* = 3 10^3^ to obtain the corresponding steady-state solutions. Various *m*-cluster solutions and their respective raster plots for different values of the network size *N* are given in Figure 3. Fig. 3(a) shows the 2-cluster solution in the network of *N* = 8, for which *m* = 2, *k* = 2, *p* = 2. In Fig. 3(b), we obtain the 1-cluster solution resulting in the synchronized behavior of all neurons in the network of *N* = 12. Also, the 2-cluster solution with *m* = 2, *k* = 2, *p* = 3 for *N* = 12 is shown in Fig. 3(c), matching the schematic diagram in Fig. 1(a). Finally, Fig. 3(d) shows the 9-cluster solution for *N* = 18, i.e., *m* = 9, *k* = 2, *p* = 1.

**Figure 3:**
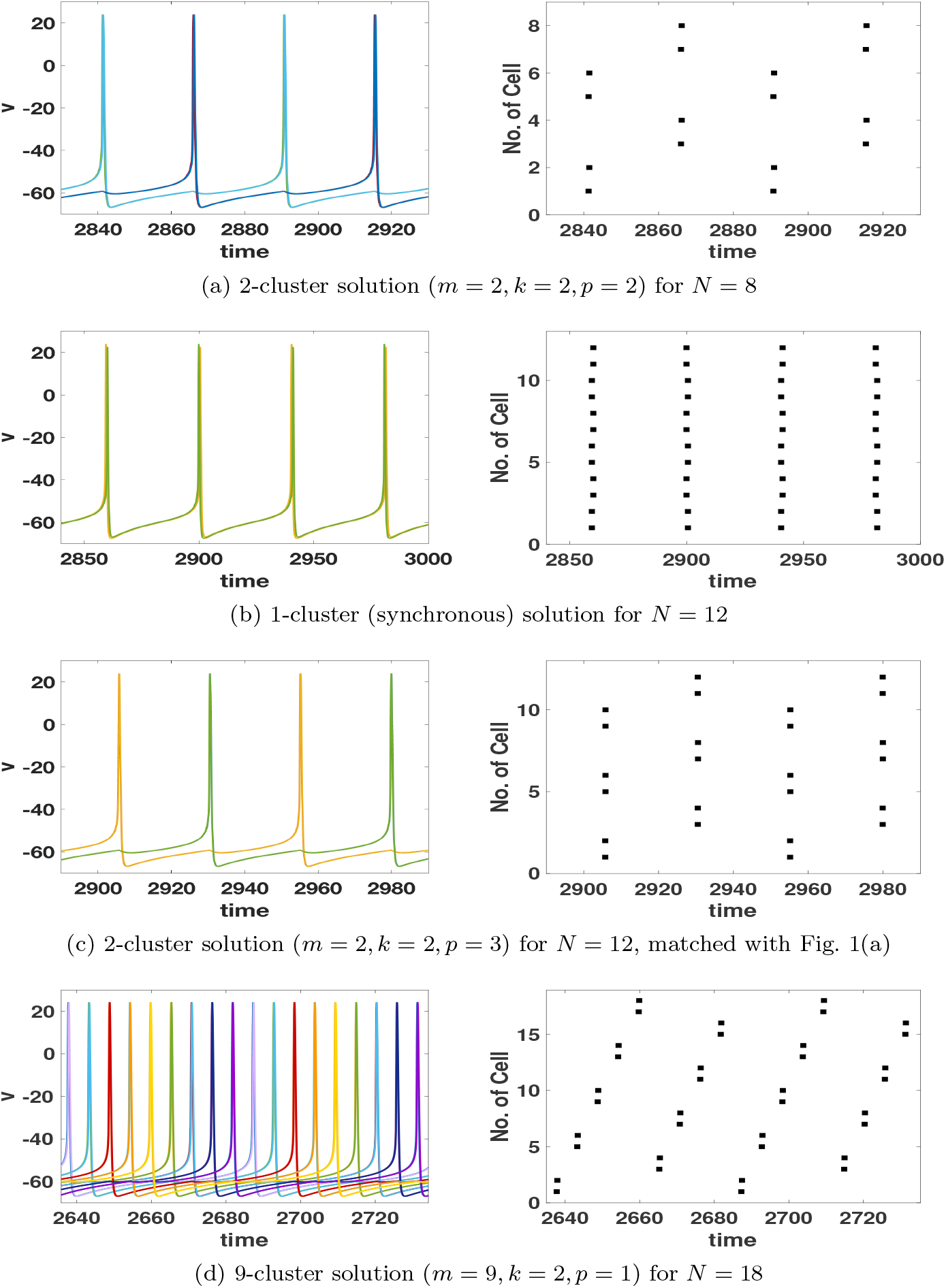
Time series (left) and its raster plot (right) for different cluster solutions with second nearest coupling only in the network for different *N* values.

We also explore the network where each neuron is connected to its third nearest neighbor only, i.e., *k* = 3. The 2-cluster solutions for *N* = 12 and *N* = 18 are demonstrated in Figs. 4(a) and 4(b), respectively. The former solution has *m* = 2, *k* = 3, *p* = 2 and the latter has *m* = 2, *k* = 3, *p* = 3.

**Figure 4:**
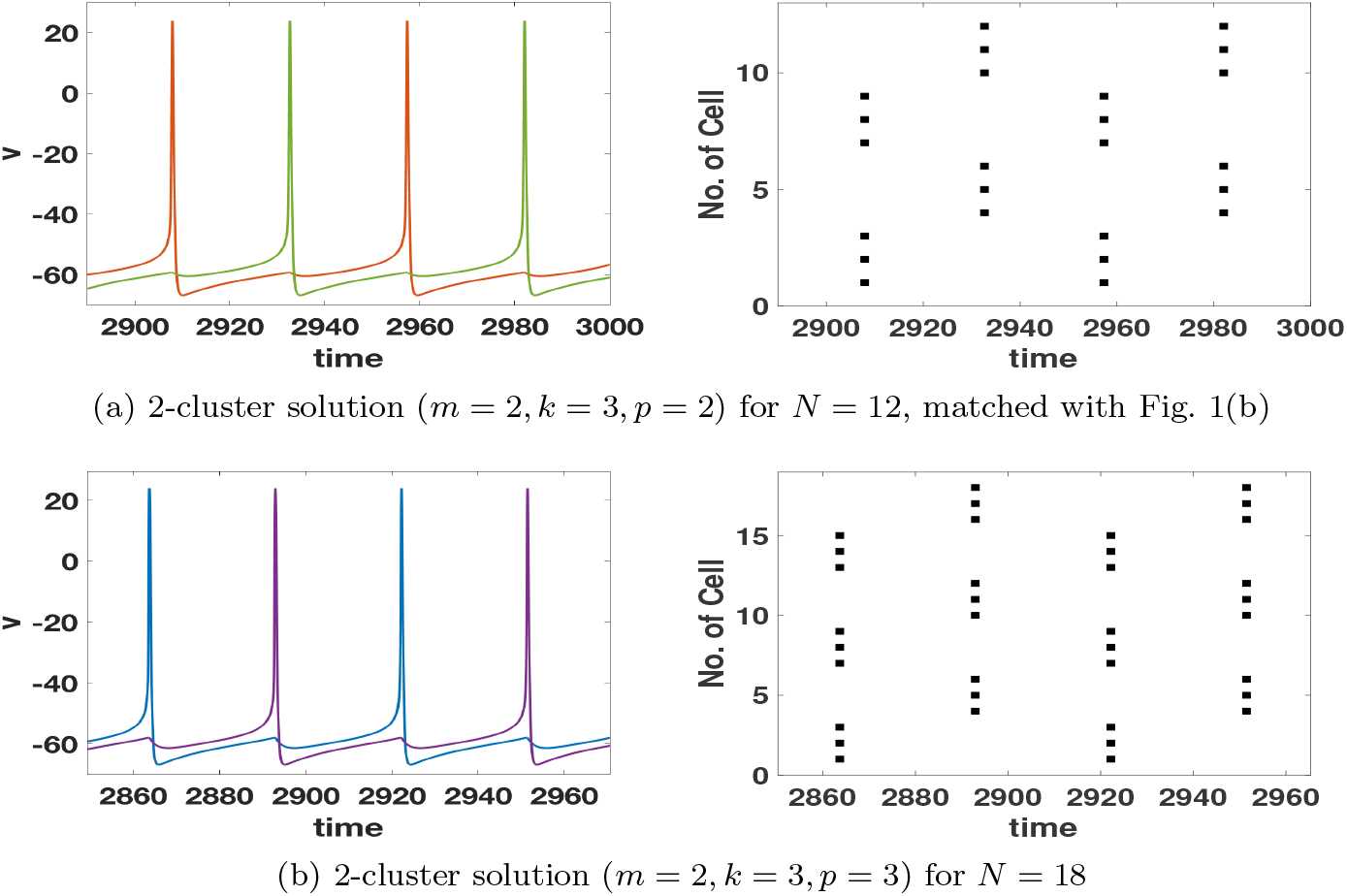
Time series (left) and its raster plot (right) for 2-cluster solutions with third nearest neighbor coupling (*k* = 3) only; (a) the network for *N* = 12 and (b) *N* = 18.

### 3.2. Non-Zero Time-Shifted Solutions

The networks considered so far have connections between every *k^th^* nearest neighbor such that each cell is not connected to its 1^*st*^, 2^*nd*^,…, (*k* - 1)^*th*^ nearest neighbors. This means that the network decomposes into *k* disjoint subnetworks (Figs. 1(a) and (b) for *N* = 12 with *k* = 2 and *k* = 3, respectively). The solutions discussed above in Sec. 3.1 focus on the case where the *k* disjoint subnetworks are synchronized, namely when cells within each cluster have the same phases (or spike at the same time) or, equivalently, have a zero time shift between them. However, other solutions, where the disjoint subnetworks are not synchronized, may exist. Such solutions would have phase differences

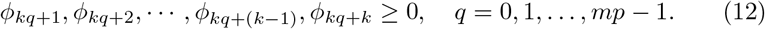

In this section, we consider these non-zero time-shifted solutions.

#### 3.2.1 Existence and stability

In the stability analysis in Sec. 3.1.1, we noted that some real parts of the eigenvalues of the circulant Jacobian matrix *A* are equal to zero. In fact, 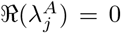 whenever *j* − 1 is a multiple of *mp*. Since 1 ≤ *j* ≤ *N* and *N* = *mkp*, there are *k* values of *j* that make the real part of the eigenvalue zero: *j* = 1, *mp* + 1, 2*mp* + 1,…, (*k* − 1)*mp* + 1.

This is related to our statement in Sec. 3.1.1 that the phase-shifts *ϕ_i_* are not independent, but satisfy the constraint 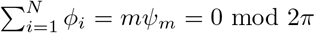. This could be used to reduce the system to fewer equations with a loss of symmetry in our system. Since our network of *N* neurons decomposes into *k* disjoint subnetworks, each with *mp* neurons, this means that 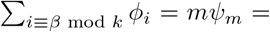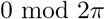 where *β* = 0,…, *k* − 1. In other words, we have *k* constraints on our system. We could reduce the system to *N* − *k* equations, but we choose not to in order to leverage the symmetry. The result is that the system will have *k* eigenvalues equal to 0 due to the *k* disjoint subnetworks. These zero eigenvalues are directly related to the existence of non-zero time-shifted solutions.

Now we show the existence of the non-zero time-shifted solution between *k* disjoint subnetworks for a network of *N* = *mkp* cells with only *k^th^* nearest neighbor coupling. The phase model for the network is described by (6) with the circulant coupling matrix

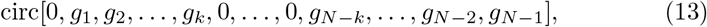

with the additional assumptions of symmetric connectivity (*g_N_*_−*i*_ = *g_i_*), *g_i_* = 0 for *i* < *k*, and *g_k_* = *O*(1) with respect to *ϵ*. Taking the phase difference *ϕ_i_* = *θ_i_*_+1_ − *θ_i_* as in Sec. 3.1.1 (1 ≤ *i* ≤ *N* and *i* is modulo *N*), we get the following system from (6) for the evolution of phase differences:

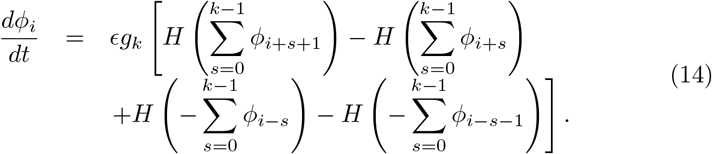

Recall that the phase differences are not independent but satisfy the constraint

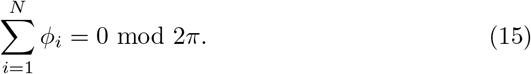

For analysis simplicity, we focus on the case of second nearest neighbor coupling (*k* = 2) only. In this case, (14) reduces to

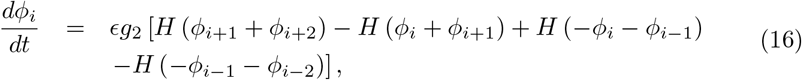

for 1 ≤ *i* ≤ *N*.

Equilibrium solutions must satisfy

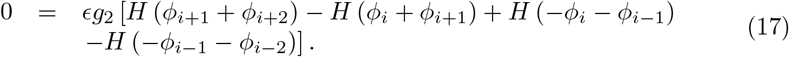

We consider solutions such that each subnetwork exhibits two clusters. In this case, cells within the same subnetwork fire in anti-phase, i.e., *θ_i_*_+2_ − *θ_i_* = *π*. Let *ϕ*_*kq*+1_ = *θ*_*kq+2*_ − *θ*_*kq*+1_ = *α* and *ϕ*_*kq*+2_ = *θ*_*kq*+3_ − *θ*_*kq*+2_ = *π* − *α* for *q* = 0, 1,…, *mp* − 1. This gives the following phase differences for arbitrary *i*:

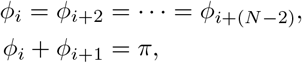

for 1 ≤ *i* ≤ *N*. Applying the above properties of *ϕ_i_* satisfies (17) as follows:

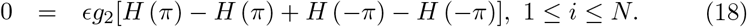

This proves the existence of non-zero time-shifted solutions for the case *k* = 2.

Note that we can easily generalize this argument to arbitrary *k*. Then, the network consists of *k* disjoint subnetworks with each subnetwork exhibiting 2-cluster firing so that *θ*_*i*+*k*_ − θ_*i*_ = *π*. This leads to the following relationships for the phase differences

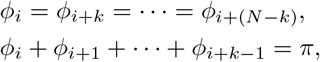

which will satisfy equilibrium solutions of (14).

Stability of these solutions stems from the stability of 2-cluster solutions in each of the disjoint subnetworks consisting of *mp* cells where *m* = 2. In this case, we can apply the stability conditions in Theorem 3.1 with *k* = 1. For example, consider the network shown in Fig. 2(a) with *N* = 12 neurons connected with their second nearest neighbors (*k* = 2) that breaks into two subnetworks of *N_sub_* = 6 neurons each. For the subnetwork of odd-numbered cells {1, 3, 5, 7, 9, 11}, we consider the stability of the 2-cluster solution: *C*_0_ = 1, 5, 9 and *C*_1_ = 3, 7, 11 where *k_sub_* = 1 and *p_sub_* = 3. Theorem 3.1 with the assumption of symmetric weights can be applied to conclude that the 2-cluster solution is locally asymptotically stable within the subnetwork with *H′_odd_*(*π*) > 0.

#### 3.2.2 Numerical results

Based on the analysis results, numerical simulations in Matlab were conducted to validate the existence and stability of non-zero time-shifted solutions using (8) and the model parameter values provided in Table 1. We simulate networks of different *N* values with second (*k* = 2) or third (*k* = 3) nearest neighbor coupling only.

We first consider the case of *k* = 2 where the network consists of two disjoint subnetworks. If the time shift between these subnetworks is nonzero, 4-cluster solutions will arise. Figure 5 demonstrates three different non-zero time-shifted solutions and their respective raster plots for the network of *N* = 8.

**Figure 5:**
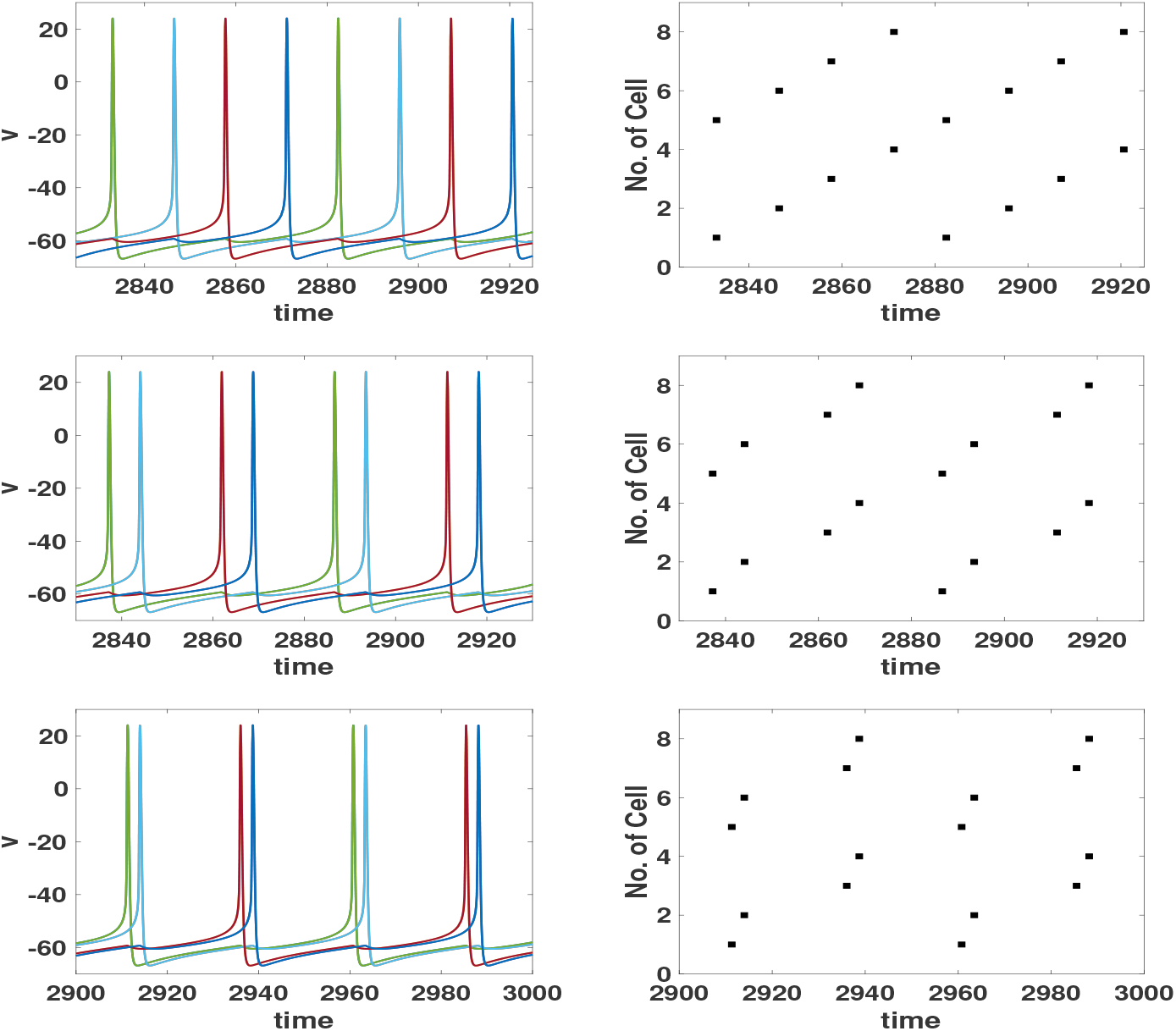
Time series (left) and its raster plot (right) for non-zero time-shifted (or 4-cluster) solutions in the network of *N* = 8 with second nearest coupling (*k* = 2) only and varying nonzero time shifts between the two disjoint subnetworks.

We also explore the network where each neuron is connected to its third nearest neighbors only, i.e., *k* = 3 for *N* = 18. In this case, the whole network consists of three subnetworks that are disjoint from each other. Each subnetwork exhibits a 2-cluster solution in which the clusters fire in anti-phase to each other. If two subnetworks fire together but the third subnetwork spikes at a different time, a 4-cluster solution arises, as shown in Fig. 6(a). On the other hand, if each of the three disjoint subnetworks fire at different times, a 6-cluster solution is found, as shown in Fig. 6(b).

**Figure 6:**
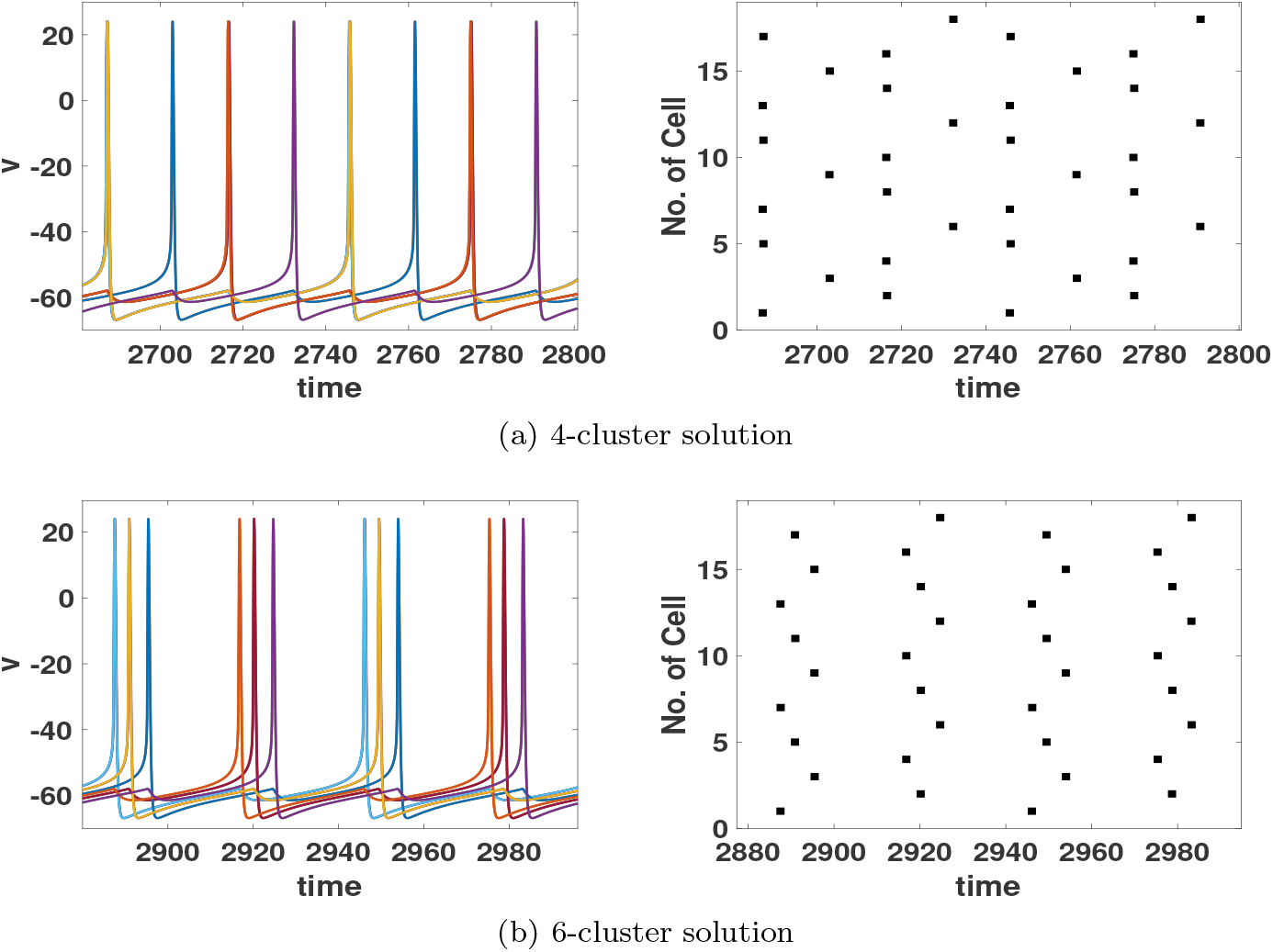
Time series (left) and its raster plot (right) for different non-zero time-shifted (or cluster) solutions in the network of *N* = 18 with third nearest neighbor coupling (*k* = 3) only.

## 4. The Second Case:*k^th^* Neighbor Coupling with Additional Couplings

In this section, we will study the effect of additional network connections on solutions studied in the previous section. The most general circulant coupling matrix with no self-coupling for the model (8) is given by

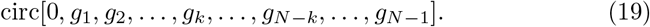

We will consider how the additional coupling terms affect the results from the previous section. We will use the phase model (6) to formulate mathematical conditions for the existence and stability of cluster solutions in the next sub-section. In the following subsection we verify our analysis through numerical simulations of the model (8).

### 4.1 Phase Model Analysis

#### 4.1.1 Existence and stability of zero time-shifted solutions

Here we consider the *m*-cluster solutions described in Section 3.1.1. We consider different possible relative strengths of the *k^th^* nearest neighbour coupling *g_k_, g_N_*_−*k*_ and the additional couplings.

The simplest situation is when *g_k_, g_N_*_−*k*_ = *O*(1) with respect to *ϵ* but the rest of the coupling strengths are *O*(*ϵ*). In this case all terms involving these *g_j_* come in at higher powers of *ϵ* in Eqs. (6) and (11). Thus we expect all the results of Theorem 3.1 to hold.

Now consider the case when all connections which exist are of equal strength, i.e., *O*(1) with respect to *ϵ*. Then we have the following.

##### Theorem 4.1.

*Consider the system (6) with N* = *mkp and coupling matrix defined by*

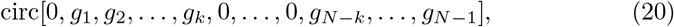

*where g_j_* = *O*(1) *with respect to ϵ. There are only two types of model-independent phase-locked solutions with adjacent neurons synchronized. The synchronous (*1*-cluster) solution exists for any choice of coupling strengths. The* 2*-cluster solution exists if N is even and the coupling strengths satisfy*

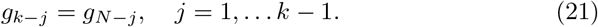

*Proof.* Using the same setup as in Section 3.1.1, define the phase difference *ϕ_i_* = *θ_i_*_+1_ −*θ_i_*, *i* = 1,…, *N*. We look for solution of the form 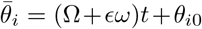 where *θ*_10_ = 0 and 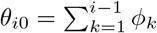 for 2 ≤ *i* ≤ *N*. We assume *N* = *mkp* and that coupling occurs between all neighbours from nearest to *k^th^*, i.e., the coupling matrix is defined by (20).

We consider an *m*-cluster solution as described by equation (9). Substituting this into equation (6) with the coupling matrix (20) gives

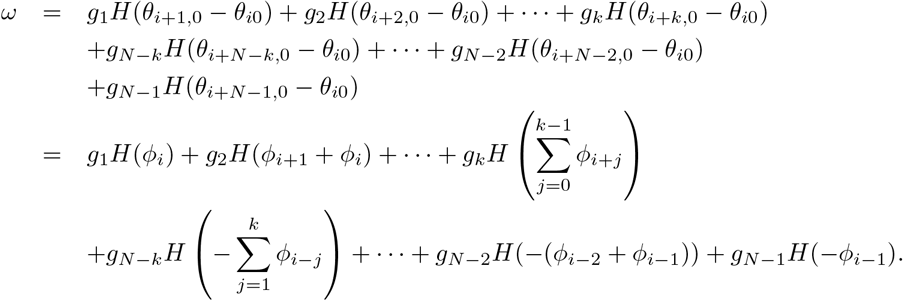

There are *k* cases to consider for the values of *ϕ_i_*, corresponding to *i* = *kq* + 1, *kq* + 2,…, *kq* + *k*. All must yield the same value of *ω* for the solution to exist. Thus we have

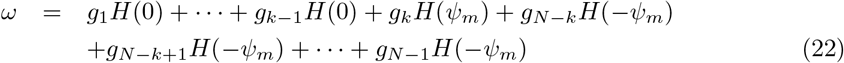

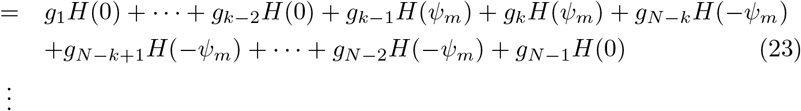

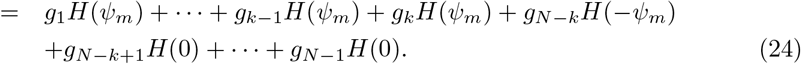

We see immediately that these equations are all satisfied for any *H* and any *g_j_* if *ψ_m_* = 0. Thus we conclude that the 1-cluster (totally synchronized) solution always exists. This is consistent with results for systems with first and second nearest neighbour coupling [31].

To proceed further, take the difference of equations (22)–(23) to find the condition

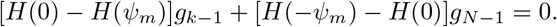

This will be satisfied for any *H* if

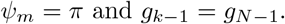

Repeating this with other pairs of equations leads to the same constraint on *ψ_m_* and the conditions (21). The result follows.

##### Remark 4.1

*It follows from the proof of Theorem 4.1 that under the conditions (21) the m-cluster solution will exist if H_odd_*(*ψ_m_*) = 0*. Thus, we do not expect any m-cluster solution with m* > 2 *to exist for every H; however, solutions for specific m may exist for a particular H. Further, other model-dependent solutions may exist under different conditions on the connection weights*.

The stability of the 2-cluster solution described in this theorem is difficult to describe for general *N*; thus we consider an in-between case, where the additional connections are weaker than *g_k_* and *g_N_*_−*k*_. We need to be careful that the weaker connections are not so weak that they are of similar strength to the neglected terms. Keeping in mind the conditions for existence of solutions derived above, we consider 2-cluster solutions (*m* = 2, *ψ_m_* = *π*) and assume the coupling matrix defined by (20) satisfies the additional condition *g*_1_*,…, g_k_*_−1_*, g_N_*_−*k*+1_,…*g_N_*_−1_ = *sδ*, while *g_k_, g_N_*_−*k*_ = *O*(1) with respect to *ϵ* and 0 < ϵ ≪ δ ≪ 1. We show two examples in Figure 7: *N* = 8 with *k* = 2 (Fig. 7(a)) and *N* = 12 with *k* = 2 (Fig. 7(b)). For the example with *N* = 12, the *O*(1) couplings are the same as in Fig. 1(a) (represented by solid lines) with weaker couplings between nearest neighbors (represented by dashed lines).

**Figure 7:**
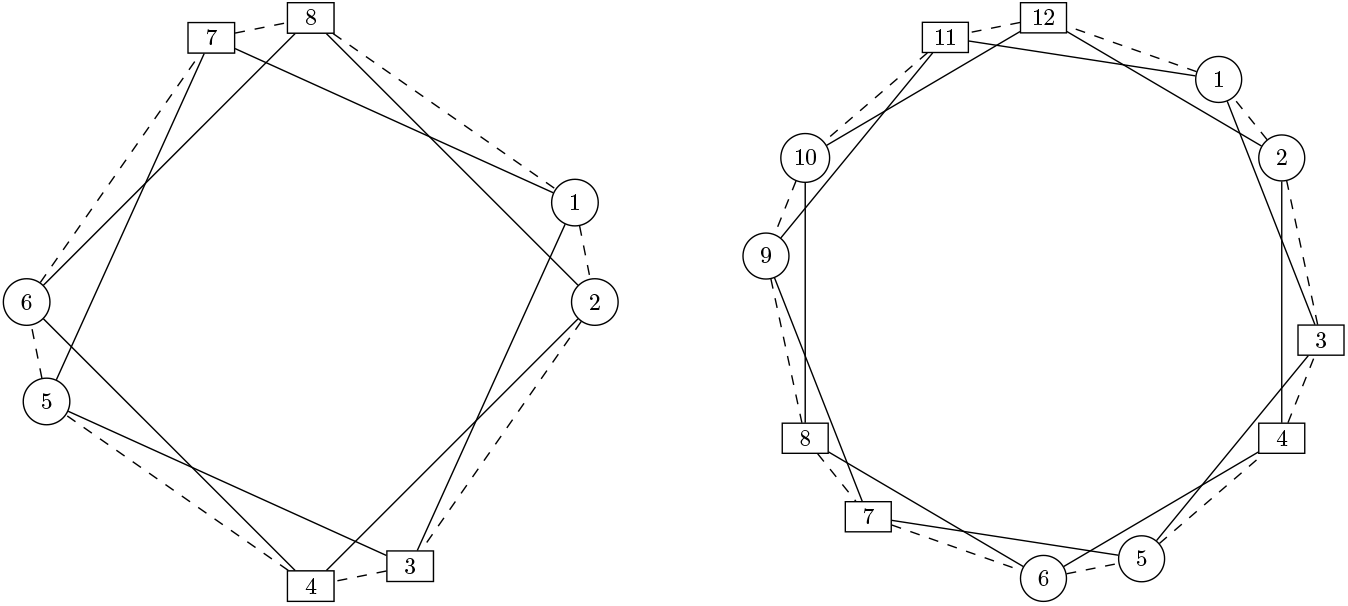
(a) Left: The network of *N* = 8 with additional coupling (in dashed lines) between nearest neighbors; (b) Right: The network of *N* = 12 with additional coupling between nearest neighbors.

We have the following result.

##### Theorem 4.2.

*Consider the model (6) with N* = 2*kp. Suppose the coupling matrix is defined by*

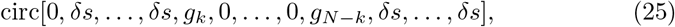

*where s, g_j_ are positive, g_k_, g_N_*_−*k*_ = *O*(1) *with respect to E and s* = *O*(1) *with respect to δ and* 0 < *ϵ ≪ δ* ≪ 1. *Then there is a* 2-*cluster solution where each cluster consists of p subgroups of k neurons. Each subgroup is synchronized and adjacent groups are π out of phase with each other. This solution is locally asymptotically stable if and only if H*′(*π*) > 0 *and H*′(0) + *H*′(*π*) > 0.

*Proof.* **Existence.** The existence of solutions follows from Theorem 4.1 since the given *N* and coupling strengths satisfy the conditions of that theorem.

**Stability.**A simple calculation shows that the Jacobian matrix for the linearization of the model (6) with the coupling matrix (25) can be written as

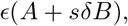

where *A* is the Jacobian matrix for the situation with no additional coupling (*δ* = 0). It follows from the analysis in Section 3.1.1 and the discussion in Section 3.2.1 that *A* has *k* zero eigenvalues and the rest of the eigenvalues are proportional (with positive constant) to −*H*′(*π*). Further, the eigenvalue 0 has geometric multiplicity *k* with linearly independent eigenvectors **v**_0_,…,**v**_*k*−1_. We take **v**_0_ = [1, 1,…,1]^*T*^.

Consideration of Eq. (6) shows that each row sum of *A* + *sδB* is 0. Thus, when *δ* ≠ 0, one zero eigenvalue persists since [*A* + *sδB*]**v**_0_ = **0**.

Now let *λ_j_* be a nonzero eigenvalue of *A* with eigenvector **v**_*j*_. Then

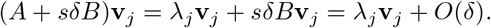

Thus, to order *δ*, *λ_j_* remains an eigenvalue for the solution. If *λ_j_* = 0, however, we have

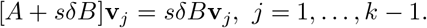

Thus the other zero eigenvalues may not persist.

For simplicity, in the rest of the proof we will take *k* = 2. The proof for other values of *k* is similar. In this case

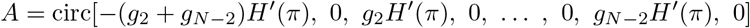

and a second, linearly independent eigenvector of the eigenvalue 0 of *A* is **v**_1_ = [1,−1, 1,−1,…,1, −1]^*T*^. Further, *B* is a banded matrix with *B_ii_* = (*H*′(0) + *H*′(*π*)) and

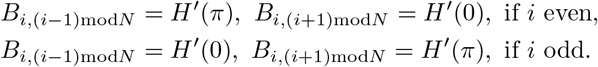

It then follows that with *δ* > 0 the second zero eigenvalue becomes −2*sδ*[*H*′(0)+ *H*′(*π*)] since [*A* + *sδB*]**v**_1_ = *sδB***v**_1_ = −2*sδ*[*H*′(0) + *H*′(*π*)]**v**_1_.

In summary, all eigenvalues except the one zero eigenvalue that persists satisfy 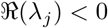 if and only if *H*′(*π*) > 0 and *H*′(0) + *H*′(*π*) > 0.

Recall that the solutions we study are of the form 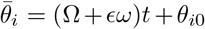. Thus the solutions correspond to lines

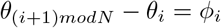

with *ϕ_i_* given by (9). The zero eigenvalue that persists corresponds to motion along these lines and hence does not affect the stability of the solutions. The result follows.

##### Remark 4.2.

*It follows from Remark 4.1, that under the conditions of Theorem 4.2 other cluster solutions may exist for* particular *models. The proof of Theorem 4.2 shows that if these solutions are unstable for the case δ* = 0*, then they will be unstable for the case δ* > 0.

#### 4.1.2 Application to the model network

For comparison with the numerical simulations below, we now consider what this analysis predicts for the model network (8) with parameters as in Table 1 and symmetric coupling matrix defined by

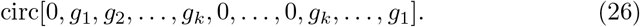

We assume that *g_k_* = *O*(1) with respect to *ϵ*.

Recall that Section 3.1.2 and specifically Table 2 gives the existence and stability for some specific values of the total number of neurons *N*, with *N* = *mkp* for some integers *m, k, p*, in the case that all couplings except *g_k_* are zero. As discussed above, if the additional couplings *g*_1_,…, *g_k_*_−1_ = *O*(*ϵ*) then these results will still hold.

Now consider the case where the additional couplings are equal and of order *δ*, *g*_1_ = *g*_2_ =…= *g_k_*_−1_ = *sδ*, where *δ* and *s* are as in Theorem 4.2. (See Figure 7 for examples with *N* = 8 or *N* = 12 and *k* = 2.) As discussed in Section 3.1.2, the synchronous solution will exist and will be asymptotically stable since *H*′(0) > 0 (see Figure 2). Further, it follows from Theorem 4.2 if *N* = 2*kp* the 2-cluster solution with *k* nearest neighbours synchronized will exist and will be stable since *H*′(*π*) > 0 and *H*′(0) > 0 from Figure 2. Finally, note that in Figure 2 we have *H_odd_*(*ϕ*) = 0 at *ϕ* ≈ *π*/3, 5*π*/3. Thus from Remark 4.1, we predict that the 6-cluster solution may exist if *N* = 6*kp* but no other *m*-cluster solutions as described in Section 3.1.1 exist. Since this solution is unstable for the case with no additional couplings (see Table 2), from Remark 4.2 we expect that it will be unstable with the additional couplings.

#### 4.1.3 Existence of non-zero time-shifted solution

In this section, we consider the existence of time-shifted solutions as described in Sec. 3.2.1 with phase differences satisfying (12) in networks with *g*_1_, *g*_2_,…, *g_k_* ≠ 0. With non-zero coupling to the first *k* nearest neighbors, the evolution equation for the phase differences *ϕ_i_* derived from (6) becomes

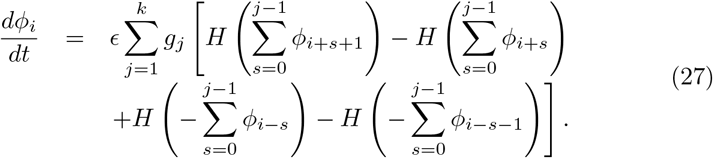

For simplicity, we consider the case *k* = 2. In this case, any equilibrium solution must satisfy the following equation:

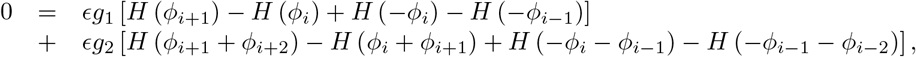

where 1 ≤ *i* ≤ *N*. From (18), we know the second term on the right-hand side is zero. Thus, for existence, the first term must be zero. Using the properties of the phase differences *ϕ_i_* given in Sec. 3.2.1 yields

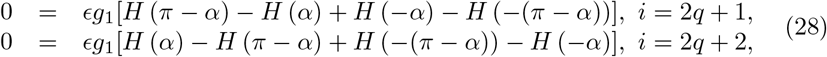

for *q* = 0, 1,…, *mp* − 1. These equations are satisfied for two conditions: i) for any time shift *α* if the function *H* is even; and ii) for *α* = *π*/2 for any *H*. For most neural network models, *H* is not an even function suggesting that the only time-shifted solution that may exist has an *α* = *π*/2 shift between cells 2*q* + 1 and 2*q* + 2 (*q* = 0, 1,…, *mp* − 1) resulting in a 4-cluster solution with *ϕ_i_* = *π*/2 for all *i*. This is a cluster solution of the type studied by [31, 7] where the neurons in a given cluster are not adjacent. Results in those papers show that this solution will be stable if and only if

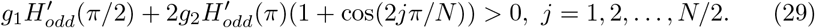

Finally, we note that since we always consider circulant coupling matrices, the system with *k^th^* nearest neighbour and additional coupling will admit *m*-cluster solutions of the type studied by [31, 7]. Conditions for the stability of these solutions can be found in those papers.

### 4.2. Numerical Results

Here, we numerically explore the existence and stability of non-zero timeshifted solutions in networks with *k* nearest neighbor coupling and additional “weak” coupling between first to (*k* − 1)^*st*^ nearest neighbors. To make the additional coupling(s) weaker than the primary coupling strength, *g_syn_* in (8), we use the scaling factor of *g_ij_* = 0.1 for |*i* − *j*| ≤ *k* − 1, *i* ≠ *j*.

In Fig. 8(a), we simulate a network of *N* = 8 neurons with second nearest coupling (*k* = 2) and weak first nearest neighbor coupling. Fig. 7(a) shows the schematic diagram of network connectivity. The network is initialized in the non-zero time-shifted solution displayed in Fig. 5(a) in which there is a time shift in firing between the two disjoint subnetworks (*ϕ*_2*q*+1_ ≠ 0, *q* = 0, 1, 2, 3) which results in a 4-cluster solution. To test its existence and stability in the presence of additional coupling(s), we set the scaling factor for the first nearest neighbor coupling strength to zero until *t* = 1, 500 ms (Fig. 8(a), top row) then set its value to 0.1 to turn on the weak additional coupling. Our results show that the non-zero time-shifted solution eventually transitions to the stable 2-cluster solution with zero time-shift (*ϕ*_2*q*+1_ = 0, *q* = 0, 1, 2, 3) (bottom row). The initial solution is close to the time-shifted solution with *ϕ*_2*q*+1_ = *α* = *π*/2. The above analysis in Sec. 4.1.3 indicates that the *ϕ*_2*q*+1_ = *α* = *π*/2 solution exists with first and second nearest neighbor coupling but based on Eq. (29) is unstable since *H′_odd_*(*π*/2) < 0 (specifically, (29) is not satisfied for *j* = 4) which agrees with our simulations.

**Figure 8:**
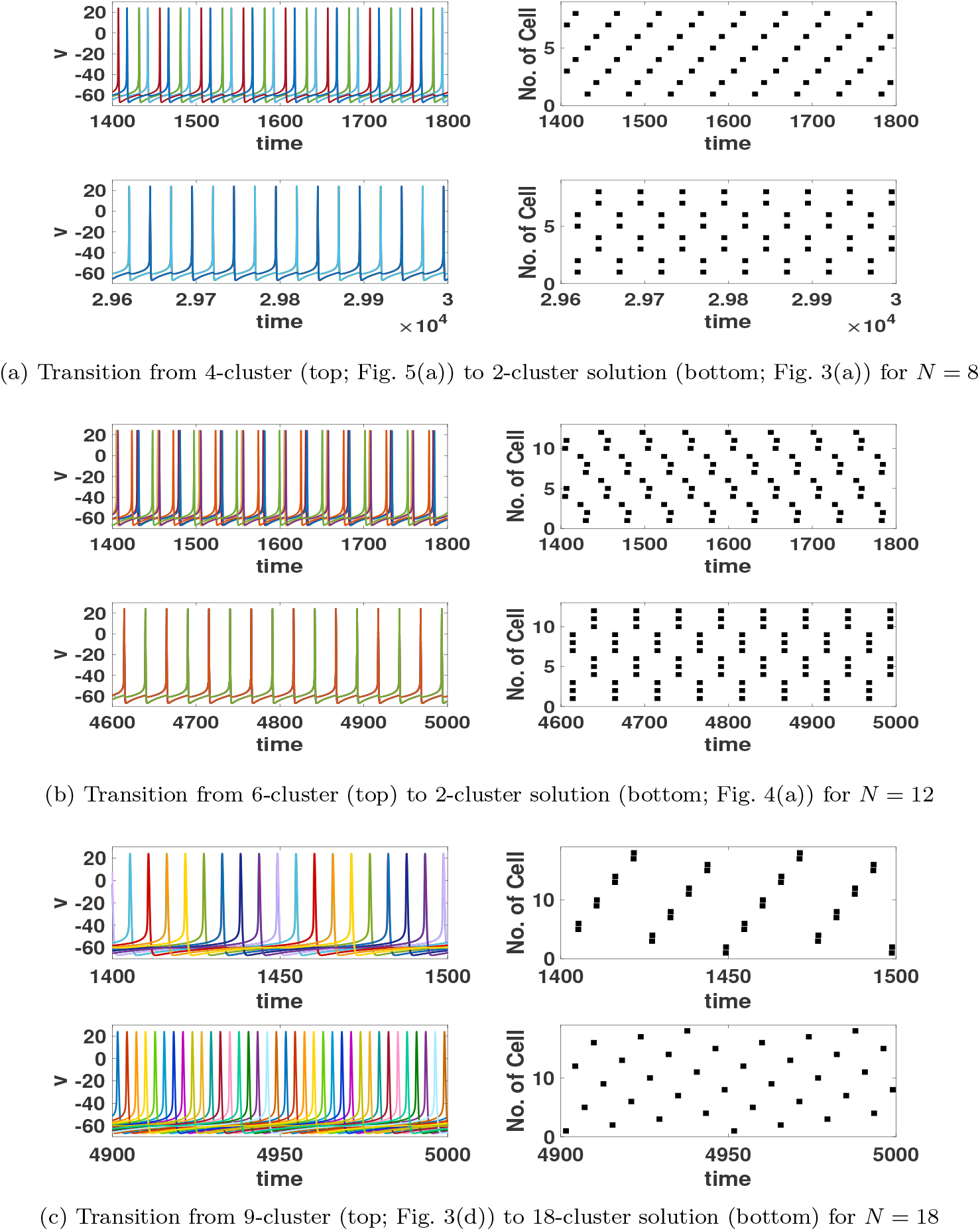
(a) and (b): Time series (left) and its raster plot (right) to show a transition from non-zero time-shifted solution to 2-cluster solution with additional coupling(s); (c): a transition from zero to non-zero time-shifted solutions with additional first nearest coupling. Additional coupling(s) were applied at *t* = 1500.

As another example, we consider a network of *N* = 12 neurons with third nearest neighbor coupling (*k* = 3) and weaker first and second nearest neighbor coupling that is initialized in a time-shifted, 6-cluster solution (Fig. 8(b), top row). Before *t* = 1, 500 ms, the scaling factor for the weaker first and second nearest coupling is zero, and then it is set to 0.1. Results show that the solution transitions to the 2-cluster solution with zero time-shift (*ϕ*_3*q*+1_ = *ϕ*_3*q*+2_ = 0, *q* = 0, 1, 2, 3) that is shown to be stable in Theorem 4.2.

Finally we show simulations that suggest that the 9-cluster solution for *N* = 18 which exists with only second nearest coupling (*k* = 2), shown in the top row of Fig. 8(c) (same as Fig. 3(d)), no longer exists when the weak first nearest neighbor coupling is added. After *t* = 1, 500 ms, when the weaker first nearest coupling is turned on, the 9-cluster solution transitions to the 18-cluster solution with equal phase difference between adjacent cells (*ϕ_i_* = *π*/18, for *i* = 1*,…,* 18, bottom row of Fig. 8(c)). These simulation results agree with the result of Theorem 4.1, which states that the only *m*-cluster solution, with adjacent neurons synchronized, that can exist with additional nearest neighbor coupling is that for *m* = 2. Since for *N* = 18 and *k* = 2 the 2-cluster solution is never possible, no *m*-cluster solution with adjacent neurons synchronized can occur. The network thus evolves to another solution type, namely the 18-cluster solution of the type studied in [31] with equal phase differences between adjacent neurons. We have verified that this solution is predicted to be stable by the phase model analysis [31, 7].

## 5. Discussion

In this paper, we study 1-D, weakly coupled inhibitory neuron networks to investigate the existence and stability of *m*-cluster solutions, in which clusters consist of groups of adjacent cells in the network that fire together. To simply describe non-monotonic, distance-dependent connectivity [38], we first consider *k^th^* nearest neighbor symmetric coupling such that each neuron in the 1-D ring is connected with its *k^th^* nearest neighbors only. This connectivity scheme results in the decomposition of the network into *k* disjoint subnetworks.

Our analysis for the *k^th^* nearest coupling only case shows that spatially localized, *m*-cluster solutions exist and can be stable. The existence of the solutions relies on the network structure rather than the network size *N*. The stability depends on the interaction function *H* and the strength of connections between cells. We use our phase model results to predict the behavior of several specific values of *N* and *k*. In particular, we can predict the number of clusters generated in the network and their stability. These results are confirmed by numerical simulations illustrating various *m*-cluster solutions for different network sizes as well as *k* values.

To study more generalized coupling, we consider the case where the network has coupling between the first to *k^th^* nearest neighbours. We show if the additional coupling (first to (*k* − 1)^*th*^) strength is sufficiently weaker than that of the *k^th^* neighbor coupling then the results with no additional coupling persist. For slightly stronger additional coupling, only two of the *m*-cluster solutions are guaranteed to persist for any model network. The synchronous (1-cluster) solution exists for any size of network and the 2-cluster solution exists if the network has an even number of neurons. We provide stability conditions for these two solutions. These results are confirmed with numerical simulations.

With only *k^th^* nearest neighbor coupling, in addition to the spatially localized, *m*-cluster solutions, our results demonstrate that solutions with more clusters exist but cells in the same cluster are spatially dispersed in the network. These solutions correspond to non-zero time-shifted solutions that emerge due to the decomposition of the network into disjoint subnetworks. In these solutions, the disjoint subnetworks are not synchronized but their cells fire with a non-zero time shift. However, when the additional coupling(s) are introduced to the network, the previously disjoint subnetworks become connected and, consequently, all but one of the non-zero time-shifted solutions no longer exist, as shown in Figure 8. The solution that retains existence displays clusters with the same phase shift between all pairs of adjacent neurons and cells in the same cluster dispersed throughout the network.

This study extends our previous work [31] in that the existence and stability conditions of *m*-cluster solutions include *k^th^* nearest neighbor coupling with asymmetric weights. Previously, we only worked with a symmetric connectivity matrix for neurons that are coupled to their first and/or second nearest neighbors on both sides. Here, we also perform a perturbation analysis to discuss the existence and stability conditions with additional coupling(s) to the *k^th^* nearest coupling. Moreover, we extend our previous work by considering nonuniform phase differences between nearest neighbors.

### Model limitations and future directions

Despite the richness of our analytical and numerical results, our results are based on the phase model reduction of an inhibitory network which explicitly assumes that neurons are weakly coupled to each other. Phase models have limitations even for studying networks of oscillatory neurons [42, 4].

Synchronization in inhibitory networks can also be obtained by mechanisms known as Interneuron Network Gamma (ING) or Pyramidal Interneuron Network Gamma (PING) [43, 24, 39, 40, 35]. In these mechanisms, synchronization of inhibitory neurons results due to gating of the timing of firing by synaptic inhibition such that cells are suppressed while inhibitory synaptic currents are active and are able to fire when synaptic currents decay. Thus, this mechanism assumes sufficiently strong synaptic coupling between cells so as to prevent firing. Our results apply to the case of weak coupling in the inhibitory network where synaptic interactions perturb the timing of cell firing without suppression of firing.

Our results focus on 1-D networks whereas the recent study [38] on nonmonotonic distance-dependent connectivity considered a 2-D network. Thus, our future study would extend the 1-D network model to investigate the 2-D network behaviors. In the 2-D setting, the diagonal neighbor coupling is a feature which could not be considered in our 1-D model. Thus we will consider its effects on cluster formation in 2-D inhibitory networks in conjunction with non-monotonic distance-dependent network connectivity.

### Conclusions

Our work may help understand cluster formation in the striatum [6, 23]. Note that the striatum has sparse, weak, unidirectional coupling [3]. Our results show that spatially localized clusters can occur when there are very few connections in the network. There has been some dispute as to whether clusters in the striatum are truly spatially compact, that is, involve only nearby neurons [23]. Our work shows that spatially compact clusters may occur, but that clusters which involve multiple groups of nearby neurons may also occur. Such solutions could correspond to clusters that involve both localized and longer range correlations between neurons as observed in [23]. Here we use reciprocal coupling in our simulations, but the mathematical results would apply to unidirectional coupling by setting some connections to zero. For example Theorem 3.1 is valid if *g_k_ >* 0 and *g_N_*_−*k*_ = 0. Since we use a general interaction function *H*, the mathematical results also apply to excitatory neurons so could be used to study cluster formation in the dentate gyrus [32].

## 6. Acknowledgement

We thank the American Institute of Mathematics for providing support through its Structured Quartet Research Ensembles program, from which this study was first initiated. SAC acknowledges the support of the Natural Sciences and Engineering Research Council of Canada.

## 7. CRediT Author Statement

All authors contributed to conceptualization, methodology, writing the original draft, reviewing and editing the final manuscript. SAC, JM, ZT and XW conducted the phase model analyses. VB and HR implemented and conducted numerical simulations.

